# Focussing on resistance to front-line drugs is the most effective way to combat the antimicrobial resistance crisis

**DOI:** 10.1101/498329

**Authors:** Meghan R. Perry, Deirdre McClean, Camille Simonet, Mark Woolhouse, Luke McNally

## Abstract

In medical, scientific and political arenas relating to antimicrobial resistance (AMR) there is currently an intense focus on multi-drug resistant pathogens that render last-line antimicrobial treatments ineffective^1–3^. We question the current emphasis of attention on resistance to last-line antimicrobials, arguing that tackling resistance to front-line antimicrobials has a greater public health benefit. Using AMR monitoring data on 25 drug-pathogen combinations from across Europe^4^, here we show that the presence of front-line pathogen resistance initiates a cascade of resistance selection that ultimately leads to pathogen resistance to last-line antimicrobials. We then interrogate, by modelling the dynamics of resistance evolution, whether 3 key interventions in the strategic response to AMR are more effectively targeted at front-line or last-line treatment. We show that interventions that make front-line therapy more effective by use of antimicrobial adjuvants or front-line resistance diagnostics or by introduction of a novel, front-line antimicrobial all lead to a larger reduction in mortality and morbidity than the same interventions implemented in last-line therapy. Mass use of a newly discovered antimicrobial in front-line infection management to maximise its public health benefit is contrary to current policy^5^ but may provide valuable incentives for drug developers. We demonstrate that funding, publications, and attention to those publications do not reflect the importance of front-line antimicrobials and are disproportionately devoted to last-line antimicrobials that account for less than 10% of antimicrobial prescriptions^6,7,8^. While studying resistance to last-line drugs is undoubtedly important, our work relays a strong message to public health agencies, funding bodies, and researchers that allocating resources to front-line infections can be a more effective way to combat the antimicrobial resistance crisis.

---

Antimicrobial resistance (AMR) is predicted over time to significantly increase rates of morbidity, mortality, and healthcare expenditure, as well as adversely affect medical advances in all clinical specialties^9^. The medical and scientific communities have reacted with a broad multi-disciplinary approach including antimicrobial stewardship programmes^10^, AMR diagnostic development^11,12^, microbiology of resistance^13,14^ and development of antimicrobial adjuvants and alternatives to antimicrobials^15^. Multi-faceted attempts are being made to incentivise and promote new antimicrobial drug development^16,17^. However, the vast scale of investment required to combat the antimicrobial resistance crisis was recently estimated to be somewhere between the Large Hadron Collider project (£6 billion) and the International Space Station ($96 billion)^18^.

In this context of limited resources interventions must be targeted to maximise reductions in morbidity and mortality, which is the ultimate aim in the fight against antimicrobial resistance. We ask if this focus should be on front-line or last-line antimicrobial management of infections? It is estimated that approximately 90% of our antimicrobial prescriptions occur in outpatient care and are for front-line antimicrobials^7,8,19^ which are the recommended first-choice for patients presenting with uncomplicated infections (for example, aminopenicillins or macrolides for community acquired pneumonias, or trimethoprim/trimethoprim-sulfamethoxazole for urinary tract infections). In addition, within the hospital there is substantial use of front-line antimicrobials particularly in the acute admissions departments. Last-line antimicrobials are administered, usually intravenously, when other antimicrobials have failed, and include agents from carbapenem, polymixin and glycopeptide classes^3^. There is a clear public health emergency with the advent of pathogens resistant to last-line agents, eg. carbapenem resistant organisms, but if we focus the predominance of our research effort on these last-line resistant organisms we argue that we are missing a valuable opportunity to minimise morbidity and mortality by effectively tackling the full clinical landscape of AMR.

What is the relationship between front-line and last-line antimicrobial resistance? It is well established that any antimicrobial use leads to increased prevalence of AMR to that antimicrobial at both the individual^20^ and the population level ^21^. Thus, the use of front-line antimicrobials is expected to lead to the evolution and spread of resistance to them. We would then expect the declining efficacy of front-line drugs to force the use of another, less commonly used antimicrobial agent, typically with a different mechanism of action. This exposure in turn increases the risk of resistance development to the second antimicrobial, and so the cascade of antimicrobial use and resistance continues, ultimately leading to pathogens resistant to last-line antimicrobials.

While this cascade of selection pressures is often implicitly assumed in the literature (sometimes conceptualised as a drug “Treadmill”^22^) there have been no tests of its occurrence. We assessed whether there is epidemiological evidence for a cascade of selection for resistance using European Centre for Disease Control antimicrobial resistance data from 2000 to 2015 for 7 pathogens (*Escherichia coli, Pseudomonas aeruginosa, Klebsiella pneumoniae, Acinetobacter spp., Enterococcus faecalis, Enterococcus faecium*, and *Streptococcus pneumoniae*)^4^ and the antimicrobials routinely used to treat them. Using mixed effects modelling, we asked how changes in levels of resistance to drug A will have future effects on levels of resistance to drugs B, C, D, etc. For example, for *E. coli*, an increase in frequency of resistance to the recommended front-line aminopenicillins will necessitate the increased use of aminoglycosides, fluoroquinolones or cephalosporins, which is reflected by an increase in the frequency of resistance to these antimicrobials in the subsequent year (Fig. 1a, Supplementary Data). For *K. pneumoniae*, which has a narrower palette of clinically effective antimicrobials, the appearance of resistance to the recommended front-line treatment with cephalosporins directly leads to increased use of the last-line drug meropenem and thus increased frequency of meropenem resistant strains in the subsequent year (Fig. 1c, Supplementary Data). Overall, our model demonstrates that increases in frequency of resistance to any antimicrobial leads to increases in frequency of strains resistant to other antimicrobials in subsequent years in 57 out of 72 (79%) inter-drug relationships (Fig. 1, Supplementary Data). Resistance in fluoroquinolones and cephalosporins in *K. pneumoniae* alone show the opposite relationship; this warrants further investigation. Overall, however, our analysis strongly supports the theory of a cascade of resistance selection from front-line to last-line antimicrobials as the general pattern of MDR evolution.

**Figure 1.**
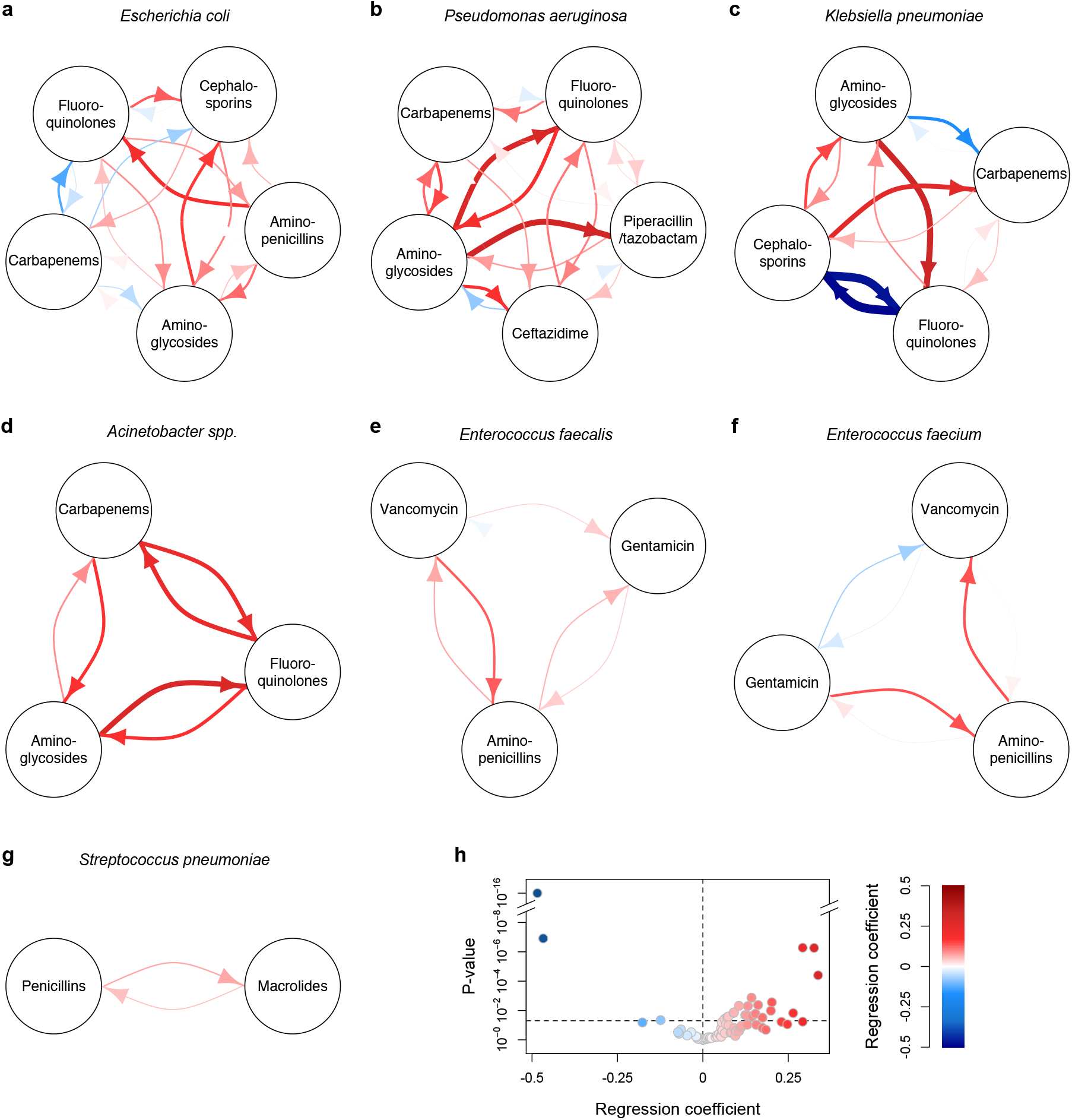
Epidemiological evidence for a cascade of resistance selection. A linear mixed effects model was applied to European Centre for Disease Control antimicrobial resistance data from 2000 to 2015. In **a-g** nodes represent resistance to different antibiotics and edges indicate the effects that frequency of resistance to an antibiotic has on frequency of resistance to another antibiotic in the subsequent year. Colours indicate the positive and negative regression coefficient and edge widths are proportional to the absolute value of the regression coefficient. For example, the red arrow from aminopenicillins to fluoroquinolones in **a** shows that, for *E. coli*, an increase in aminoglycoside resistance in the current year will lead to an increase in fluoroquinolone resistance in the subsequent year. **h** summarises the regression coefficients and p-values for all inter-drug effects in **a-g**.

Given that last-line drug resistance evolves via a cascade of selection following the appearance of resistance to front-line drugs where should we focus our intervention efforts to maximise public health benefit? We hypothesised that focussing our efforts on front-line interventions that optimise efficient clearance of pathogens resistant to front-line drugs will ultimately have the largest public health benefit. The three interventions that we have examined are key areas in the strategic response to AMR: 1) introduction of antimicrobial adjuvants that maximise efficacy of existing antimicrobials against resistant strains, 2) introduction of point-of-care resistance diagnostics, and 3) introduction of a brand-new antimicrobial with a novel mechanism of action.

To test our front-line hypothesis, using a susceptible-infected compartmental model we model the dynamics of resistance evolution in a focal pathogen species with four possible strains - a susceptible strain, a strain resistant to a front-line treatment, a strain resistant to a last-line treatment, and a multi-drug resistant (MDR) strain that is resistant to both treatments (Fig. 2a). Our population comprises of two classes; a ‘carrier’ class of individuals that are colonised with the pathogen but do not develop disease from that focal pathogen and a ‘symptomatic’ class of individuals who develop disease with this pathogen and thus require treatment. In this ‘symptomatic’ class, we consider a scenario where patients are initially treated with the front-line drug, before escalation to the last-line treatment if their infection does not clear. The clearance and onward transmission of each strain within and between carrier and symptomatic classes is affected by susceptibility to the treatment and the cost of resistance mutations, and we allow for the *de novo* evolution of resistance during treatment. Individuals within the ‘carrier’ class can be colonised with any strain. Since individuals in this class do not develop disease due to our focal pathogen, they do not receive treatment for our focal pathogen. However, carriers can receive antimicrobial treament for an infection with a different pathogen, or owing to inappropriate antimicrobial usage (e.g. self-medication or use of an antibiotic to treat a viral infection). This exposure to antimicrobials while in the pathogen is in carriage and not the target of treatment is referred to as “bystander selection”^19^. We let the level of resistance evolve in this population for a specified burn-in time period and then evaluated the benefit of the intervention for a period of time after the intervention. We used four different measures of public health outcomes. At the population level, we assessed the total disease induced mortality after intervention, and the average prevalence of infection (as a measure of disease induced morbidity). At the individual patient level we assessed the probability that a patient dies of their infection once infected, and also the average duration of infection as a measure of morbidity.

**Figure 2:**
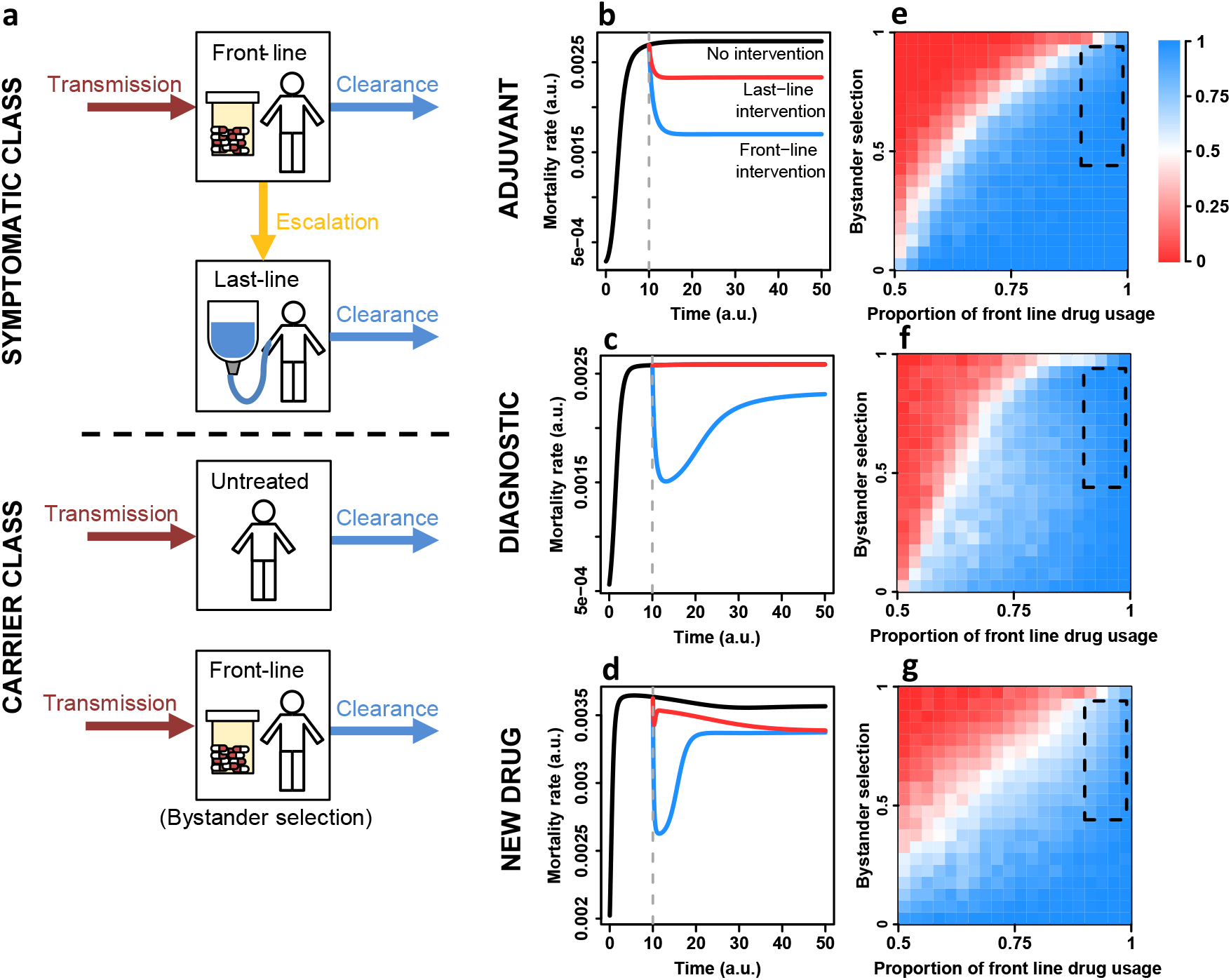
Front-line interventions lead to lower mortality than last-line interventions. Panel **a** shows a schematic of the structure of our model. Panels **b-d** show examples of the observed dynamics of instantaneous mortality rate over time in the absence of intervention (black line), under front-line intervention (blue line) or under last-line intervention (red line) for the three types of interventions (adjuvant, diagnostic, new drug). The intervention is introduced after an initial “burn-in phase” where AMR evolves in the absence of intervention (dashed vertical line). Panels **e-g** show the proportion of simulations in which front-line intervention led to lower mortality than last-line intervention as a function of bystander selection and the proportion of front-line drug usage. The colour map goes toward blue when the front-line intervention led to a lower total mortality in more than 50% of the simulations (front-line intervention optimal), towards red for less than 50% of the simulations (last-line intervention optimal). The black-dashed box defines the area of this parameter space corresponding to current antimicrobial usage practices, based on estimates of bystander selection and proportion of front-line drug usage (see Methods).

We first examine the effects of a hypothetical antimicrobial adjuvant that partially restores the efficacy of a drug against resistant pathogens. This could, for example, represent optimisation of drug delivery to increase an effective dose against a resistant strain^23^ or addition of molecules targeting resistance mechanisms, e.g. novel beta-lactamase inhibitors or novel CRISPR phage therapeutics^15,18,24^. We find that focussing on optimising front-line treatment (i.e. with adjuvants to increase the clearance of front-line resistant strains) decreases the basic reproductive number (*R*_0_) of MDR strains to a greater extent than last-line intervention (Supplementary Fig. 1), and leads to better public health outcomes (Fig. 2, Supplementary Fig. 2). This is particularly evident in the demarcated area that represents current antimicrobial usage patterns, where front-line drugs account for 90% to 97% of antimicrobial prescriptions and the percentage of antimicrobial exposures that constitute bystander selection ranges from 44% to 94% (Fig. 2, see Methods). The reason for this result is simple: while interventions targeting last-line resistance only have a direct effect on selection for last-line resistance, if we improve the efficacy of a front-line treatment against resistant infections then patients’ pathogenic organisms will mainly be cleared without the need to use last-line treatment. Thus, development of a front-line adjuvant as an intervention decreases morbidity, mortality, and the *R*_0_ of MDR pathogens both directly by clearing infections early and indirectly via reducing last-line drug usage.

In assessing a hypothetical point-of-care resistance diagnostic intervention, many of which are under development^25^, we consider a scenario where a diagnostic would either detect resistance to a front-line or a last-line antimicrobial. With the front-line diagnostic intervention, if resistance to a front-line drug is detected patients are directly treated with a last-line antimicrobial. With the last-line diagnostic intervention, if last-line resistance is detected, patients do not escalate to last-line treatment. The results of this diagnostic model again show greater decreases in mortality and morbidity when diagnostics are applied at the front-line. A front-line resistance diagnostic detects the need for last-line treatment immediately without patients having to progress through failure of front-line treatment and therefore receive effective treatment earlier. This therefore leads to more rapid recovery and lower morbidity and mortality (Figure 2, Supplementary Fig. 2). This model output matches previous clinical studies which demonstrate the positive impact of prompt appropriate antimicrobial therapy on morbidity and mortality^26^. We note however, that in the absence of other interventions both front-line and last-line diagnostics have no effect on the fitness of MDR strains – there is no advantage to knowing that these strains are resistant when there is no alternative drug to which they are sensitive (Supplementary Fig. 1)^27^.

Finally, to assess where in the treatment hierarchy to deploy a newly developed antimicrobial we extend our model to three different antimicrobials. We consider two scenarios: 1) reservation of the new antimicrobial as a last-line drug (as is current policy^5^) and 2) deployment of the new antimicrobial as a new front-line drug. The model shows clearly here that deployment of a newly discovered drug as a front-line treatment leads to greater reductions in mortality and morbidity than saving this drug for last-line therapy (Figure 2, Supplementary Fig. 2). It leads to greater reductions in the fitness of front-line resistant, last-line resistant and MDR strains (Supplementary Fig. 1). This reduction is largely as a result of increasing the clearance of strains resistant to the previous front-line treatment by using a drug with a novel mechanism of action. This increases the efficacy of the first treatment a patient receives, shortening the length of infection, giving less opportunity to transmit, and leading to less patients requiring escalation to middle and last-line drugs to clear their infection. This is in contrast to reserving the novel antimicrobial for last-line usage, where patients may have already received treatment with front and middle line antimicrobials to which their infections were resistant.

Resistance can and will ultimately develop to the new drug as by introducing it as either a front or last-line treatment we have simply added a new step in the resistance cascade. However, deployment of the new antimicrobial as a front-line drug has a much greater initial benefit before resistance to this drug emerges and spreads, resulting cumulatively in lower morbidity and mortality. This result, as with the other two interventions, is robust to the choice of length of burn-in phase before intervention and the length of post-intervention evaluation period (Supplementary Figures 3–6). We have proved our results analytically for a simplified model without the carrier class of hosts (Supplementary Equations).

If a novel antimicrobial is discovered it is a topic of considerable interest as to the most effective way for it to be incorporated into the clinical landscape. Drug companies are arguing currently that there is no financial incentive for them to discover a new antimicrobial that will only be used sparingly because it is being preserved carefully for patients who have MDR pathogens^5^. Our work clearly shows that there is a strong public health advantage to a new drug being introduced in front-line settings, as it ultimately will lead to the largest reductions in morbidity and mortality. Promised bulk use of a new agent is far more economically attractive to drug companies and would help to promote novel antimicrobial research and development to provide the healthy drug pipeline required.

Our models clearly demonstrate the benefit of front-line interventions but we show below that the research community is focussing disproportionately on last-line resistance. Hitherto, there has been no published data on how much research effort in response to the AMR crisis is allocated to front-line or last-line antimicrobials. We addressed this information gap by combining data for 13 antimicrobial classes on worldwide antimicrobial usage^6^ with six different measures of research focus. We assessed the level of funding research into resistance to each antibiotic class using both total grant dollars awarded and total numbers of active grants from the National Institute for Health (NIH), the world’s largest public biomedical funder^28^. We quantified the current focus of research groups using numbers of research papers published since 2012 focusing on resistance to each antibiotic class. Finally, we used three different measures of the attention these papers received using Altmetric, a service that tracks the online sharing and coverage of research^29^. In all cases, we find that there is no correlation between research focus on different antibiotic classes and their usage, with infrequently used last-line drugs receiving a disproportionate amount more funding, published research articles, and attention to those articles compared to expectation based on their usage (Fig. 3, Supplementary Fig. 7, Supplementary Table 1). The extent of this disproportionate focus is extreme. Penicillins constitute 38% of global antimicrobial usage, while carbapenems constitute just 0.2% of usage, yet research into carbapenem resistance received more funding than penicillin resistance. The seven least commonly used classes of antimicrobials (Chloramphenicols, Aminoglycosides, Carbapenems, Rifamycin, Glycopeptides, Monobactams, and Polymixins) constitute only 3% of global usage yet received 47% of research funding. This pattern is also not explained by newer antimicrobial classes, where it is likely that less is known about resistance mechanisms, receiving more attention as we found no relationship between the age of an antimicrobial class and any attention metrics (Supplementary Figure 8, Supplementary Table 2).

**Figure 3.**
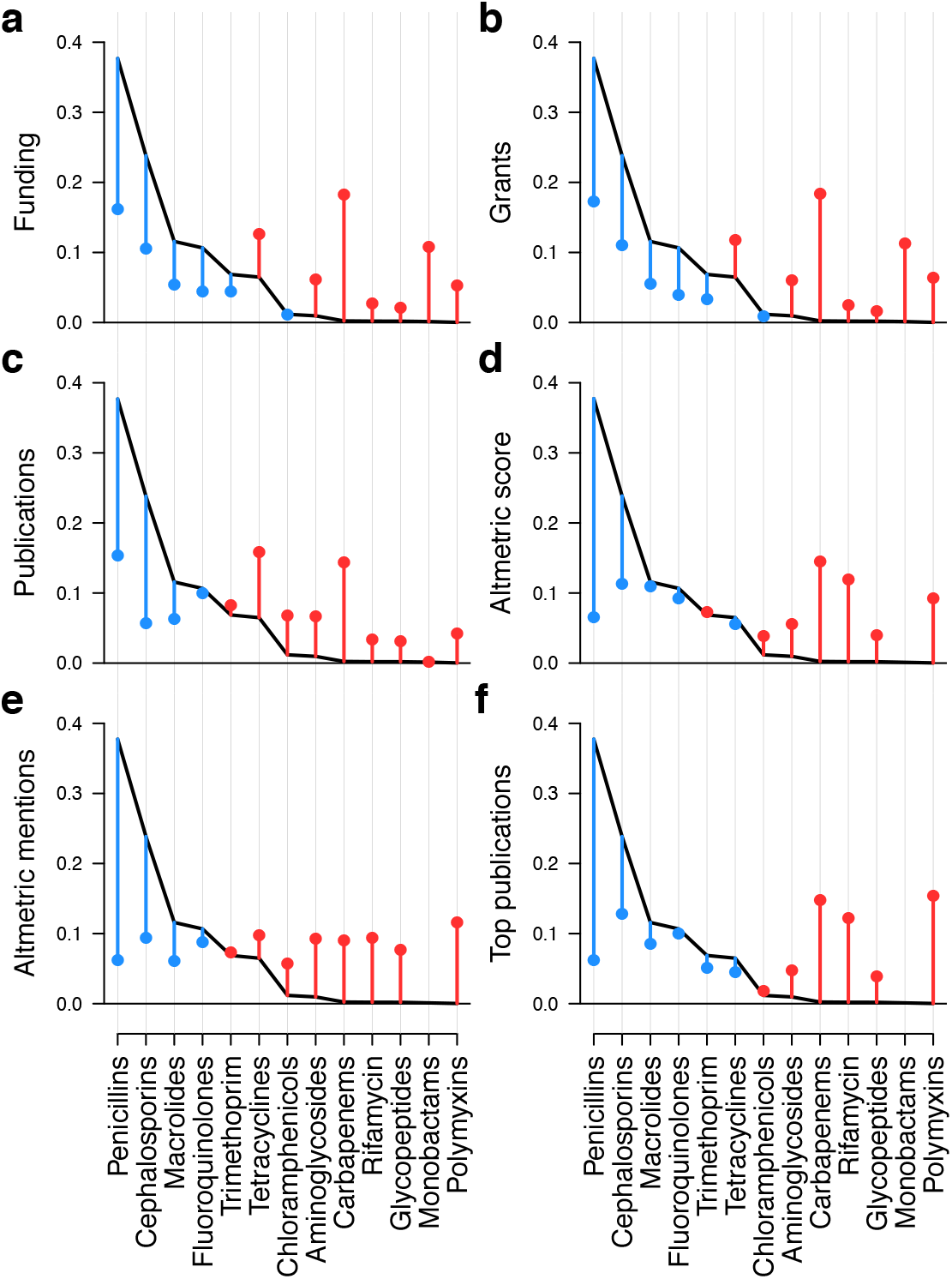
Resistance to rarely used last-line antimicrobials receives a disproportionate amount of research attention given their consumption. Points show the share of each attention metric (i.e. proportion of total) that each resistance to each class of antimicrobials receives. The black line shows the proportion of total global antimicrobial consumption that antimicrobial class is responsible for. Points are coloured red if the proportion of attention for resistance to that class exceeds its proportion of consumption, and blue if it is less than that proportion of consumption. The attention metrics are **a**, NIH funding dollars, **b** NIH grants, **c** publications, **d** Altmetric score, **e** Altmetric mentions, and **f** contribution to top 10% of publications by Altmetric score. In all instances more commonly used front-line antimicrobials receives proportionally less attention than expected based on their usage, while resistance to the least commonly used antimicrobials receive proportionally more attention than expected.

Our analysis of current research practices demonstrates that current research focuses disproportionately on resistance to last-line antimicrobials: we are focusing on the end of the resistance cascade, rather than the beginning. The fundamentals of medicine and epidemiology teach us that such an approach may be misguided – it is generally better to tackle a disease process in its early stages to prevent future clinical events, which carry substantial morbidity. Our epidemiological analysis and mathematical modelling show that the old adage ‘look after the pennies and the pounds will look after themselves’ can be applied; if the treatment of front-line resistant pathogens is optimised this has a greater public health impact than focussing on last-line resistant pathogens.

The occurrence of MDR pathogens that are resistant to last-line antimicrobials such as carbapenems is a public health emergency and requires attention. However, as a medical and research community we need to act in a way that will minimise the ongoing impact of AMR and use our resources and any newly discovered antimicrobials to achieve the most substantial decreases in morbidity and mortality. Our results show that the way to utilise resources most effectively is to urgently re-evaluate our priorities and increase our focus on ways to tackle resistance to front-line antimicrobials.

## Acknowledgements

We thank Kristofer Wollein Waldetoft, Meriem El Karoui and Sam Brown for comments and discussion, and Christine Tedijanto for sharing estimates of bystander selection. M.R.P. was supported by the Institutional Strategic Support Fund ISSF J22738, L.M. was supported by a Chancellor’s Fellowship from the University of Edinburgh, and M.W. acknowledges support from the ‘Selection and Transmission of Antimicrobial Resistance in Complex Systems [STARCS]’ project in the Joint Programming Initiative on Antimicrobial Resistance (grant no. MR/R000093/1). L.M. and M.W. also acknowledge support from The Novo Nordisk Foundation (NNF16OC0021856: Global Surveillance of Antimicrobial Resistance).

## Author contributions

M.R.P. and L.M. conceived the project, D.M. collected the data, D.M. and L.M. analysed the data, C.S and L.M. performed the mathematical modelling with input from M.W. and M.R.P., and M.R.P. and L.M. drafted the paper. All authors contributed to conceptual development, interpretation of results, and manuscript revision.

---

**Supplementary Figure 1.**
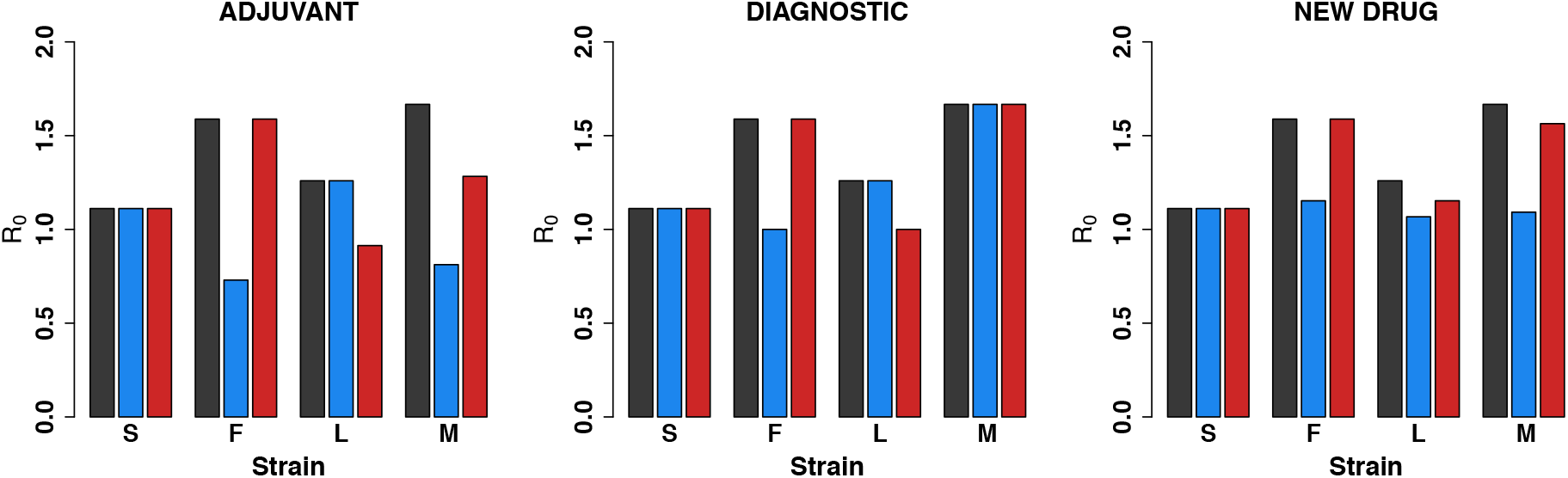
Front-line interventions are more effective at reducing pathogen *R*_0_. Shown are the *R*_0_ values for each strain (S = sensitive, F = front-line resistant, L = last-line resistant, M = multi-drug resistant) under no intervention (black bars), front-line intervention (blue bars), and last-line intervention (red bars). For the “adjuvant” model front-line intervention reduces the *R*_0_ of strains F and M to a greater extent than last-line intervention. Last-line intervention has a greater impact on strain L, but this strain is less likely to be common as front-line drugs will be used more that last-line drugs. A similar result is seen for the “new drug” model. For the diagnostic model front-line intervention has a greater impact on strain F, while last-line intervention has a greater impact on strain L, though again strain F is likely to be more common as the front-line drug is used more. Diagnostic intervention does not impact strain M in the absence of other intervention as there are no effective drugs to treat this strain. All *R*_0_ values are calculated using the simplified model without carriage.

**Supplementary Figure 2.**
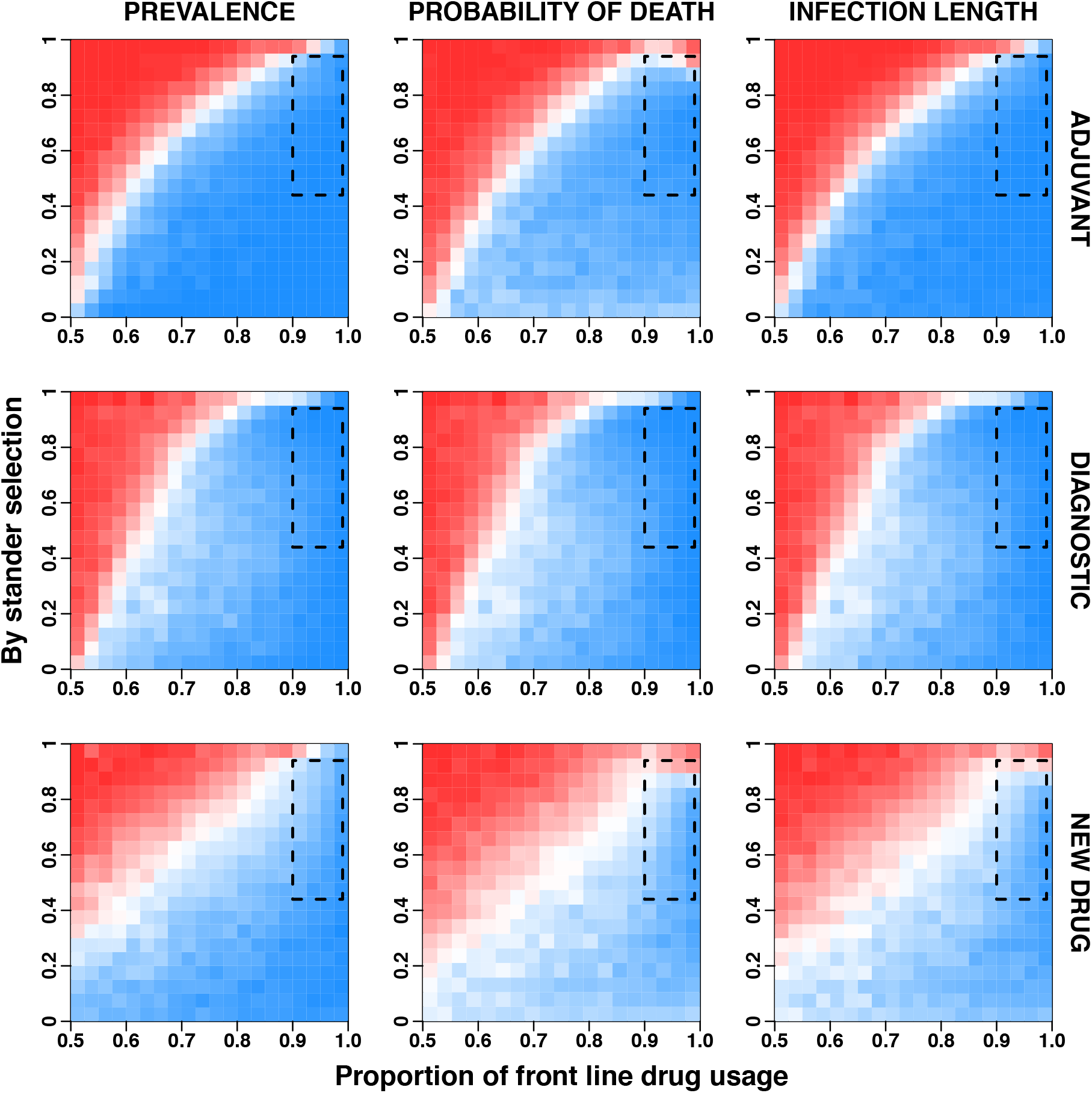
Front-line intervention is superior for other public health measures. The plot is as in Fig. 2e-g, but columns here are for three different public health measures (prevalence of infection, probability that a patient dies of infection, and average infection length). For all measures front-line intervention is superior in the relevant region of parameter space.

**Supplementary Figure 3.**
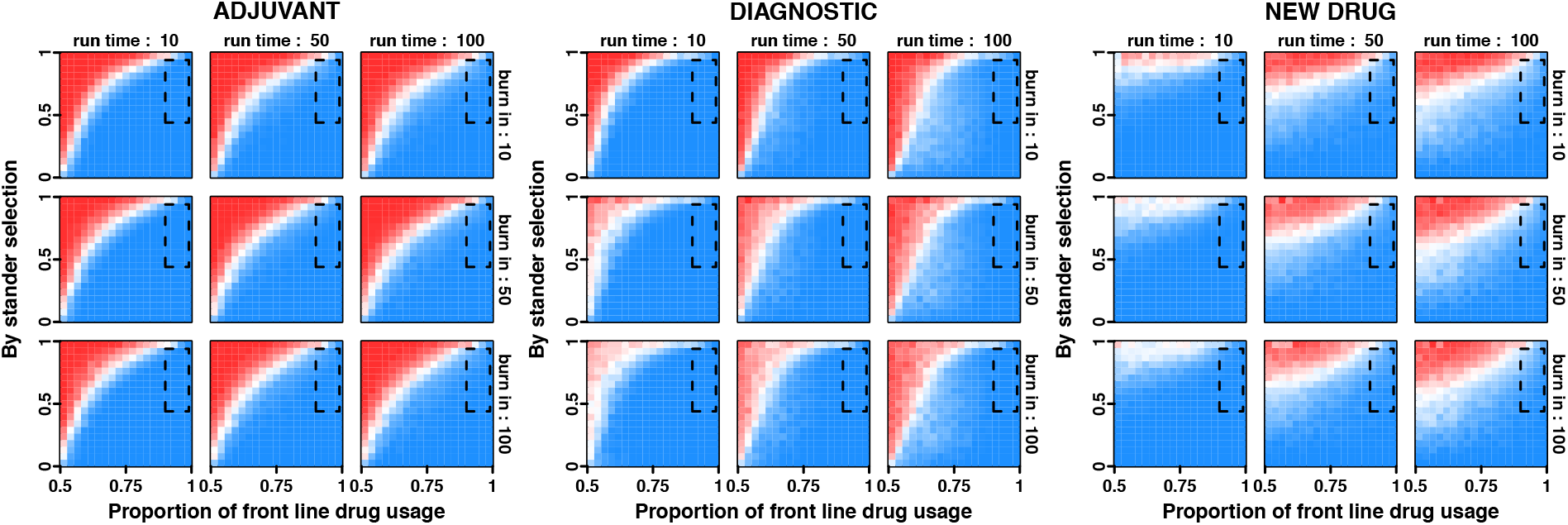
Front-line intervention leads to a lower total mortality regardless of burn-in period or run time. The plot is as in Fig. 2e-g, but showing results for different simulation burn-in periods and run times (total simulation length) for total mortality as an outcome measure. In all cases front-line intervention leads to lower total mortality in the relevant region of parameter space.

**Supplementary Figure 4.**
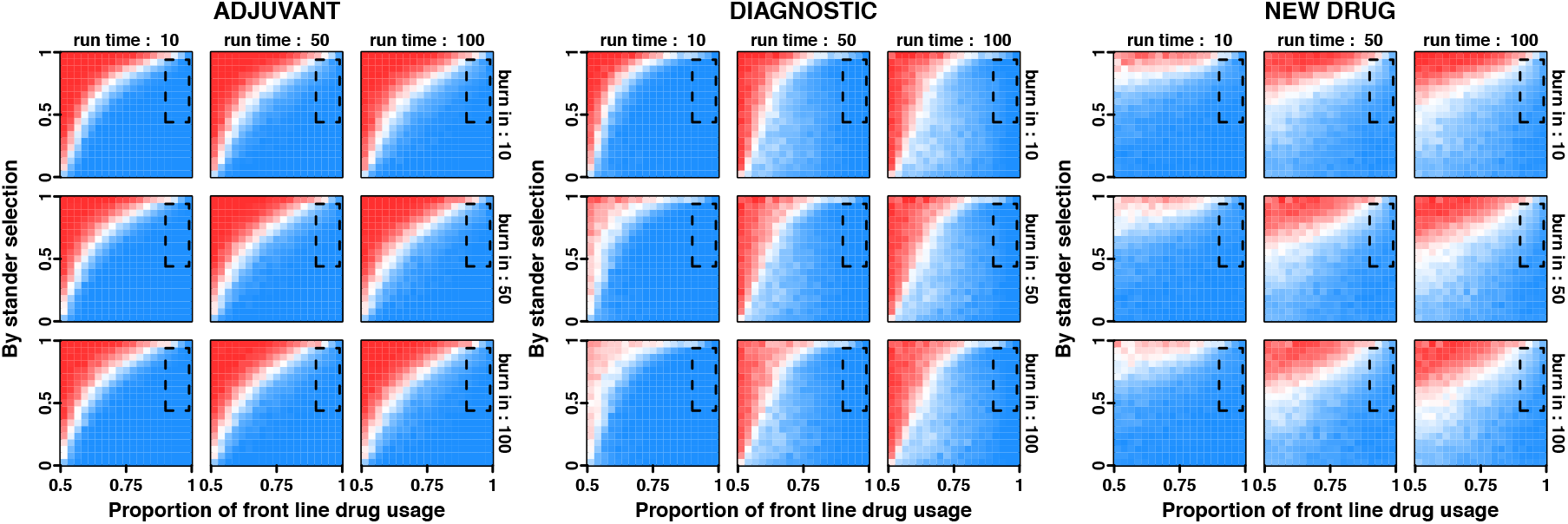
Front-line intervention leads to a lower prevalence of infection regardless of burn-in period or run time. The plot is as in Fig. 2e-g, but showing results for different simulation burn-in periods and run times (total simulation length) for the prevalence of infection as an outcome measure. In all cases front-line intervention leads to lower prevalence of infection in the relevant region of parameter space.

**Supplementary Figure 5.**
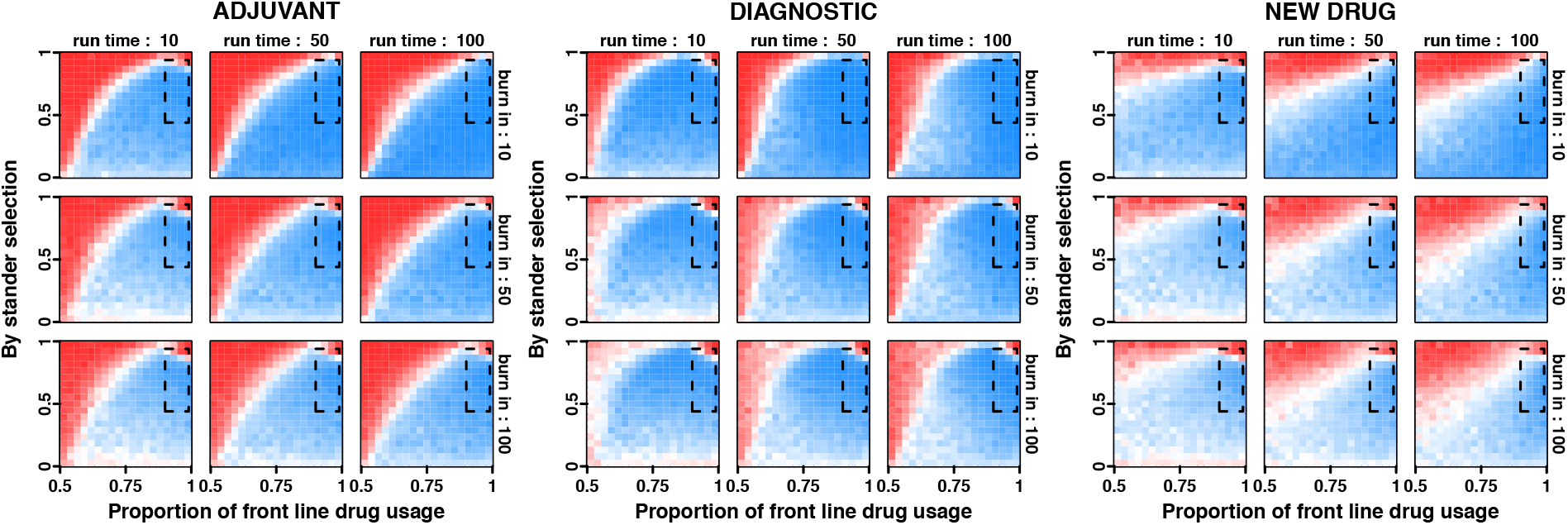
Front-line intervention leads to a lower probability of death regardless of burn-in period or run time. The plot is as in Fig. 2e-g, but showing results for different simulation burn-in periods and run times (total simulation length) for the probability that a patient dies from infection as an outcome measure. In all cases front-line intervention leads to lower prevalence of infection in the relevant region of parameter space.

**Supplementary Figure 6.**
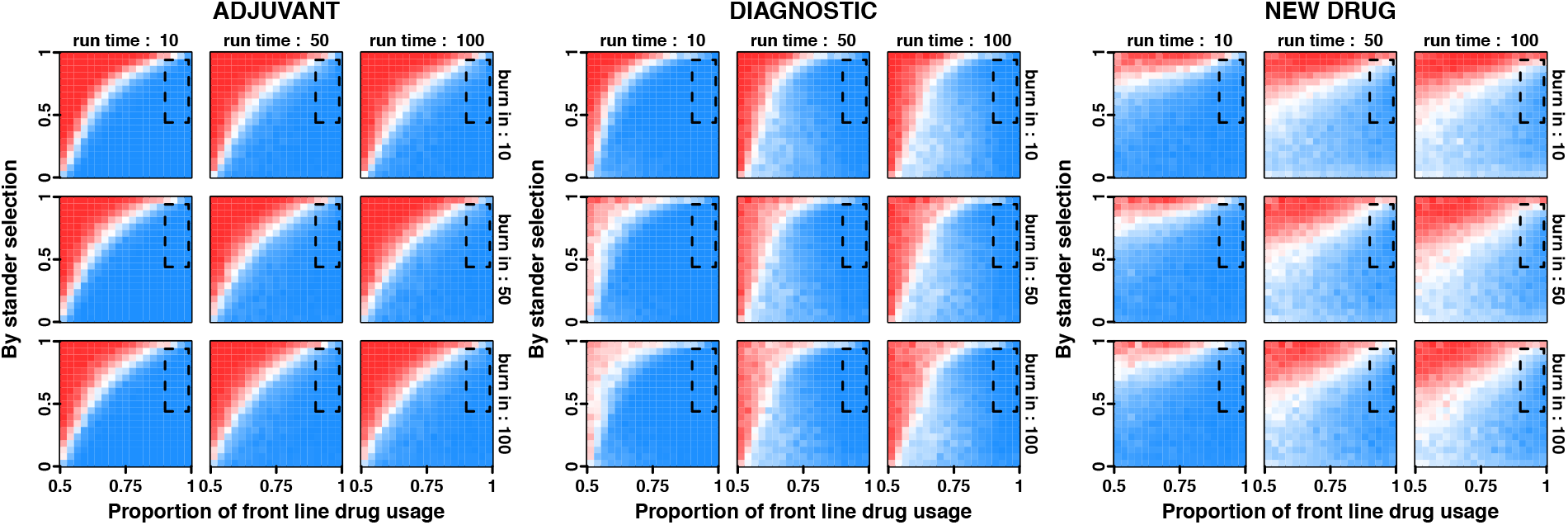
Front-line intervention leads to a lower average length of infection regardless of burn-in period or run time. The plot is as in Fig. 2e-g, but showing results for different simulation burn-in periods and run times (total simulation length) for the average length of infection as an outcome measure. In all cases front-line intervention leads to lower prevalence of infection in the relevant region of parameter space.

**Supplementary Figure 7.**
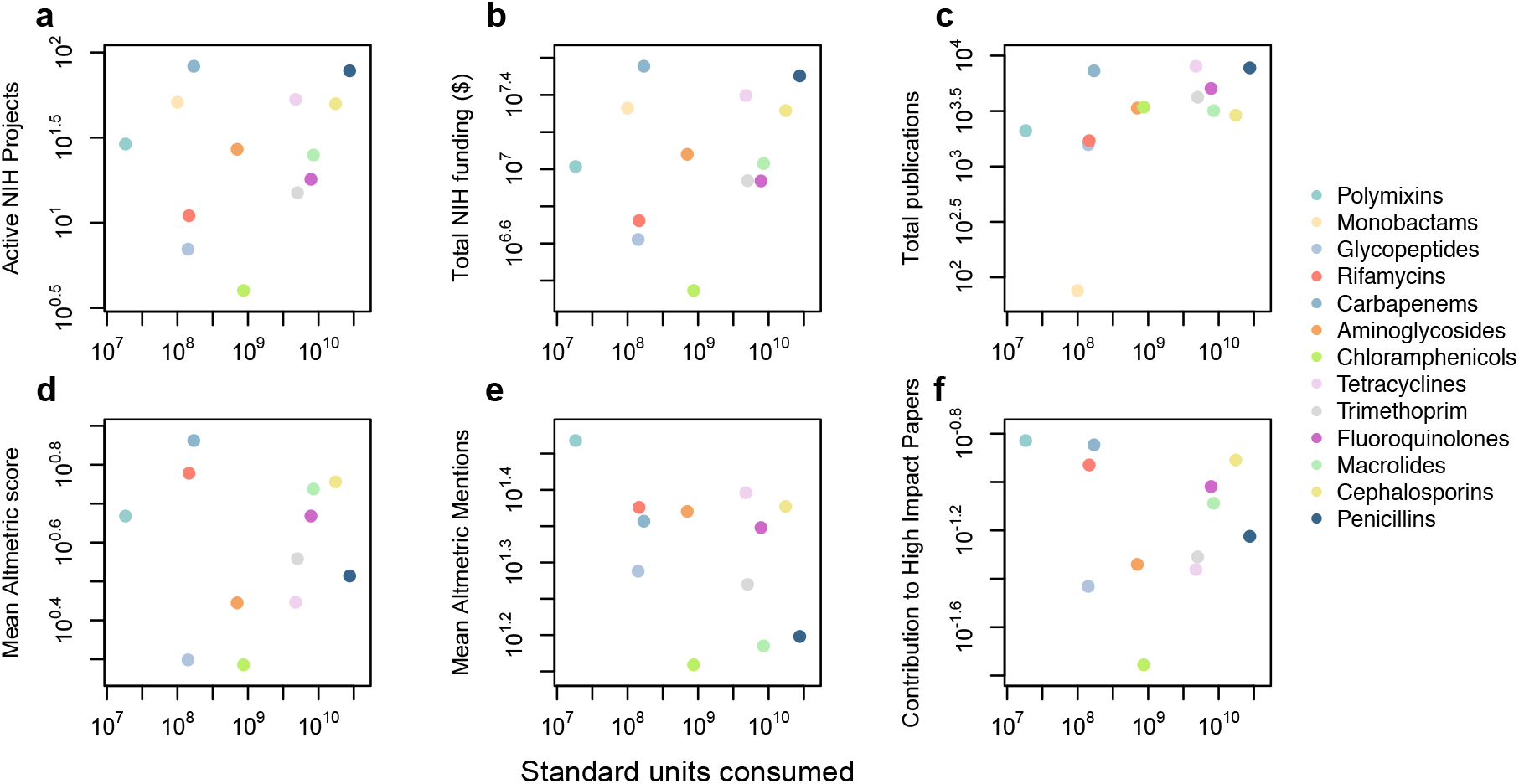
There is no correlation between research focus and antimicrobial usage. Plotted are 6 log_10_-transformed measures of research focus (**a** active NIH grants, **b** total NIH funding, **c** publications since 2012, **d** mean Altemetric score, **e** mean Altmetric mentions, and **f** contribution to top 10% of papers by altmetric score) against log_10_-transformed worldwide antimicrobial consumption. Colours indicate the class of antimicrobial. In all cases there is no significant relationship between consumption and research attention.

**Supplementary Figure 8.**
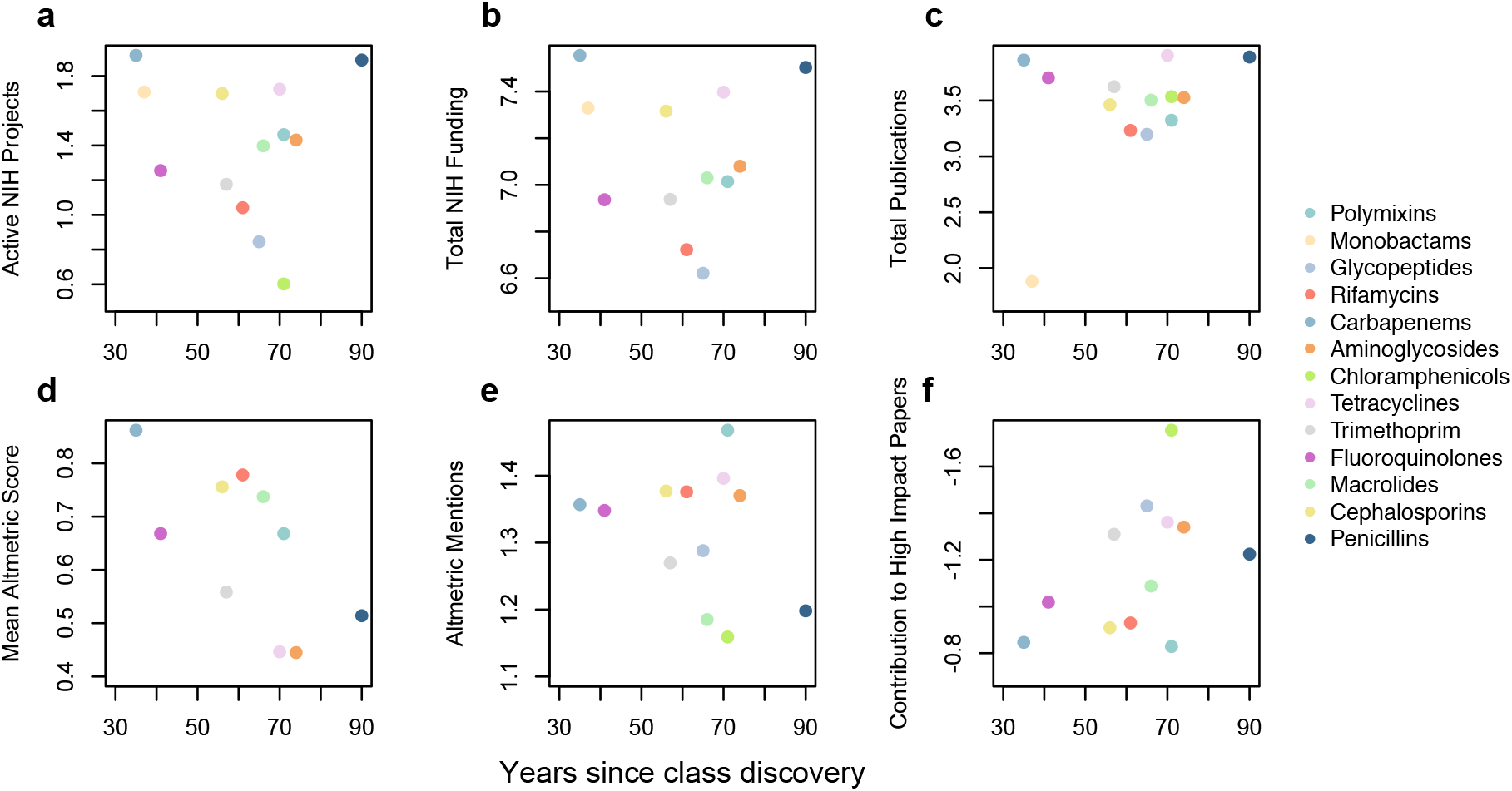
There is no correlation between research focus and the age of an antimicrobial class. Plotted are 6 log_10_-transformed measures of research focus (**a** active NIH grants, **b** total NIH funding, **c** publications since 2012, **d** mean Altemetric score, **e** mean Altmetric mentions, and **f** contribution to top 10% of papers by altmetric score) against the number of years elapsed since discovery of the antimicrobial class. Colours indicate the class of antimicrobial. In all cases there is no significant relationship between antimicrobial class age and research attention.

**Supplementary Table 1.** Relationship between global antibiotic usage for 13 classes of antibiotics and measures of research focus. depicted in Fig. 3 and Supplementary Fig. 7. Shown are slope and intercept estimates from our regression models with associated standard error, t-values and p-values for both standard regression models and those looking at deviation from the predicted relationships, and R-squared values.

| Consumption |  | Estimate | Std. Error | t-value | P- value | t-value as deviation from predicted | P-value as deviation from predicted | R squared |
| --- | --- | --- | --- | --- | --- | --- | --- | --- |
| **Total NIH Active Projects** | *Intercept* | 0.65 | 1.07 | 0.6 | 0.56 | 8.29 | < 0.001 | 0.043 |
|  | *Slope* | 0.08 | 0.12 | 0.7 | 0.5 | -7.8 | < 0.001 |  |
| **Total NIH Funding** | *Intercept* | 6.26 | 0.94 | 6.659 | < 0.001 | 9.39 | < 0.001 | 0.063 |
|  | *Slope* | 0.089 | 0.104 | 0.856 | 0.41 | -8.81 | < 0.001 |  |
| **Sum of Papers post 2012** | *Intercept* | 1.066 | 1.218 | 0.875 | 0.4001 | 5.93 | < 0.001 | 0.26 |
|  | *Slope* | 0.262 | 0.134 | 0.875 | 0.298 | -5.51 | < 0.001 |  |
| **Mean Altmetric Score** | *Intercept* | 0.602 | 0.548 | 1.098 | 0.0764 | 17.81 | < 0.001 | 0.0001 |
|  | *Slope* | -0.002 | 0.06 | -0.032 | 0.975 | -16.76 | < 0.001 |  |
| **Mean Altmetric Mentions** | *Intercept* | 1.715 | 0.246 | 6.971 | < 0.001 | 41.36 | < 0.001 | 0.2099 |
|  | *Slope* | -0.044 | 0.027 | -1.63 | 0.134 | -38.92 | < 0.001 |  |
| **Contribution to High Impact Papers** | *Intercept* | -0.736 | 0.803 | -0.916 | 0.381 | 12.63 | < 0.001 | 0.02866 |
|  | *Slope* | -0.0476 | 0.088 | -0.543 | 0.599 | -11.96 | < 0.001 |  |

**Supplementary Table 2.** Relationship between global antibiotic usage for 13 classes of antibiotics and age of antibiotic class. depicted in Fig. 3 and Supplementary Fig. 8. Shown are slope and intercept estimates from our regression models with associated standard error, t-values, p- and R squared values.

| Years Since Discovery |  | Estimate | Std. Error | t-value | P- value | R squared |
| --- | --- | --- | --- | --- | --- | --- |
| **Total NIH Active Projects** | *Intercept* | 1.61 | 0.477 | 3.373 | **0.006** | 0.019 |
|  | *Slope* | -0.0035 | 0.0076 | -0.461 | 0.65 |  |
| **Total NIH Funding** | *Intercept* | 7.283 | 0.423 | 17.225 | **< 0.001** | 0.026 |
|  | *Slope* | -0.0036 | 0.0067 | -0.541 | 0.599 |  |
| **Sum of Papers post 2012** | *Intercept* | 2.67 | 0.577 | 4.63 | **< 0.001** | 0.145 |
|  | *Slope* | 0.013 | 0.0092 | 1.364 | 0.599 |  |
| **Mean Altmetric Score** | *Intercept* | 1.0397 | 0.2179 | 4.773 | **< 0.001** | 0.314 |
|  | *Slope* | -0.007 | 0.0034 | -2.14 | 0.058 |  |
| **Mean Altmetric Mentions** | *Intercept* | 1.438 | 0.1268 | 11.345 | **< 0.001** | 0.089 |
|  | *Slope* | -0.0019 | 0.00196 | -0.986 | 0.058 |  |
| **Contribution to High Impact Papers** | *Intercept* | -0.6233 | 0.3486 | -1.788 | 0.104 | 0.206 |
|  | *Slope* | -0.0087 | 0.0054 | -1.608 | 0.139 |  |

## Methods

### Analysis of ECDC data on frequency of resistance

To assess whether resistance to other drugs results in increases in resistance to a focal drug we used data from the ECDC’s European Antimicrobial Resistance Surveillance Network (EARS-Net). This database includes data on the percentage isolates resistance to a range of antibiotics for 7 pathogen species (*Escherichia coli, Pseudomonas aeruginosa, Klebsiella pneumoniae, Acinetobacter spp*., *Enterococcus faecalis*, *Enterococcus faecium*, *Streptococcus pneumoniae*, and *Staphylococcus aureus*) for 30 European countries from 2000 to 2015 (though some pathogens are only available more recently) and is available at http://atlas.ecdc.europa.eu. We omitted *Staphylococcus aureus* from our analysis as only data on methicillin resistance are available, so it cannot be used to assess effects of levels of resistance of different drugs on a focal drug’s resistance dynamics. All resistance data (percentage of isolates resistant) were log-odds transformed prior to analysis to homogenise variance. For each antibiotic and pathogen for which data were available we fit a random effects model of the following form

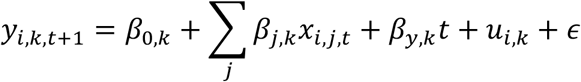

where *y_i,k,t+1_* is the change in the log-odds of resistance (first difference) between year *t* and *t* + 1 in country *i* for antibiotic *k, β_0,k_* is the intercept, *x_i,j,t−1_* is the level of resistance in country *i* for antibiotic *j* in year *t, β_j,k_* is the effect of resistance level for antibiotic *j* on the change in resistance for the focal antibiotic, *β_y,k_* is a trend in the change in resistance frequency, *u_i_* is a random effect of country, and *ϵ* is the normally distributed error term. All models were fitted in R using the lme4 package. Full model results are given in Supplementary Data.

### Mathematical model of intervention consequences

In order to assess whether interventions to tackle last-line or front-line drug resistance would have the greatest public health benefit we modelled the dynamics of resistance evolution using a susceptible-infected compartmental model. We considered a scenario where patients initially receive a front-line antibiotic and then transition to last-line antibiotic treatment at a constant rate if they remain infected. All infections are cleared at a baseline rate, with additional clearance rate when treated with an antibiotic to which the infecting strain is susceptible. Ill patients die from infection at a constant rate. Each resistance mutation increases a strain’s clearance rate, representing the cost of resistance. Each strain also transmits to uninfected individuals at equal rates. Patients can be infected with four different strains, a susceptible strain *S*, a strain resistant to the front-line drug *F*, a strain resistant to the last-line drug *L*, and a multi-drug resistant strain carrying both resistance mutations *M*. We further assume that some individuals carry the infection asymptomatically (“carriers”). The pathogen may still experience exposure to antimicrobial usage when in carriage either owing to its use to treat other infections or because of inappropriate usage (e.g. use of an antibiotic to treat viral infections). This exposure to the antimicrobial when not causing symptoms is termed “bystander selection”. We assume that the focal pathogen only experiences bystander selection for the front-line drug as there are no current estimates of bystander selection for last-line drugs and this assumption works against our hypothesis by increasing selection for resistance to front-line drugs. We considered three possible interventions: introduction of an adjuvant that blocks the resistance mechanism, introduction of a diagnostic to detect resistance and avoid use of the antimicrobial when resistance is present, and introduction of a novel antimicrobial drug. We evaluated the effects of each intervention in terms of four public health measures: total mortality, total morbidity, probability of dying upon infection, and average duration of infection. For full details of the model, it’s numerical implementation, and mathematical proofs of our results see the Supplementary Equations.

To assess which region of parameter space corresponds to current antimicrobial usage practices we assessed two key quantities, the proportion of drug usage that is front-line and the proportion of antimicrobial exposures that are bystander selection. For the proportion of drug usage that is front-line we took 0.9 as the lower bound as >90% of antimicrobial prescriptions are outpatient and front-line^7,8,19^, and an upper bound of 0.97 as the six most commonly used classes of antimicrobials (penicillins, cephalosporins, macrolides, fluoroquinolones, trimethoprim, and tetracyclines), all of which are considered front-line, constitute 97% of global usage. For bystander selection we used the estimates of ref. 20 of the proportion of drug exposures that constitute bystander selection for 9 pathogens, which were calculated using patterns of outpatient antimicrobial prescribing, infection incidence, and pathogen carriage in the US. We took a weighted mean of bystander selection values across drugs for each pathogen, weighing by the proportion of antimicrobial prescriptions targeting that pathogen that each drug represents. This yielded estimates of proportions of antimicrobial prescriptions that are bystander selection for each pathogen of 0.86 for *Streptococcus pneumoniae*, 0.94 for *Haemophilus influenzae*, 0.85 for *Moraxella catarrhalis*, 0.9 for *Staphylococcus aureus*, 0.76 for *Escherichia coli*, 0.79 for *Pseudomonas aeruginosa*, 0.9 for *Klebsiella pneumoniae*, 0.89 for *Streptococcus* agalactiae, and 0.44 for *Streptococcus pyogenes*. We took the range of these values (0.44 – 0.94) as the realistic range of bystander selection proportions.

### Assessment of current AMR research focus

We gathered data on 13 different drug classes (Tetracyclines, Chloramphenicols, Penicillins, Cephalosporins, Monobactams, Carbapenems, Trimethoprim, Macrolides, Aminoglycosides, Fluoroquinolones, Glycopeptides, Polymyxins, and Rifamycin) in order to assess the current focus of AMR research.

We use the antibiotic consumption data from Van Boeckel *et al* 2014^6^, see therein for full methods. These data are gathered from the IMS Health MIDAS database (IMS Health, Danbury, CT, USA). These data estimate antibiotic consumption from the volume of antibiotics sold in both hospital and retail pharmacies from numerous countries (a total of 63 used here) across the globe (see Supplementary Table 3 in Van Boeckel *et al* 2014). The data used are given in Standard Units derived from an algorithm using regional, sector and distribution channel factors to project national estimates of antibiotic consumption for the years 2000 and 2010. The table provides information for 16 drug classes. For our analysis we summed the data from the three classes of Penicillins and omitted Carbacephems due the paucity of data available from Altmetric, publication and funding sources from our analysis, leaving us with 13 drug classes.

We use RePORTER data from the U.S. Department of Health and Human Services, National Institutes of Health (NIH)^28^ to get estimates for total number of active grants (those that have not yet reached the end of their budget period) and funding amount awarded for resistance to each of the 13 classes of antibiotic. The data are drawn from a number of databases (eRA databases, Medline, PubMed Central, the NIH Intramural Database, and iEdison) and provide information on projects funded by NIH, Centers for Disease Control and Prevention (CDC), Agency for Healthcare Research and Quality (AHRQ), Health Resources and Services Administration (HRSA), Administration for Children and Family (ACF), and U.S. Department of Veterans Affairs (VA). For our analysis we used the Text Search function of the RePORTER database for projects with *“drug name* AND resistance”.

To assess the total numbers of papers published for each of our drug classes we used the Web of Science database to search all databases for each of our drug classes with ‘*resistan*\*’ in the title, abstract and keywords of papers across all years. In figure 1 we used total number papers for each drug class published from 2012 onwards to align with the Altmetric scores, which were only available from 2012. To ensure that this choice of cut-off did not effect our results we analysed numbers of publications using only papers publishes post 2012, only papers published pre 2012, and papers published in any year, all of which showed comparable results (Supplementary Table 1).

To measure the impact of publications on the classes of antibiotics we used the Altmetric Explorer database^29^ to gather information on the mean Altmetric scores, mean number of online mentions for each of our antibiotic classes with ‘resistance’, and the probability contribution of each drug class resistance to the top 10% scores of high impact research papers. Altmetrics are data encompassing the number of mentions of the search terms across a number of sources including mainstream media coverage (mentions on news sites for example), Faculty of 1000, research blog mentions, citations on Wikipedia and in public policy documents, bookmarks on reference managers like Mendeley, and mentions on social networks such as Twitter and Facebook, which link to the published article since 2012. The Altmetric scores are calculated based on the number of mentions and weighted by source to account for projected reach of the outlet. This means that while the majority of mentions come from Twitter, a mention on Twitter is given a weighting of “1” while a mention on a news site might be given a score of “8”. Together these provide information about how articles are discussed online across the globe. Monobactams were omitted from the analysis for Altmetric scores due to lack of data leaving us with 12 classes of antibiotic for this analysis.

For each of these metrics of research focus we first assessed whether there was a correlation between research attention and antimicrobial consumption using linear models of log_10_ transformed research attention and antimicrobial consumption data, and found no significant correlation for any research attention metrics (Supplementary Fig. 7, Supplementary Table 1). We then also tested if the relationship between drug consumption and the measure of research focus deviated from that expected if research focus was proportional to drug consumption. This expected relationship is given by

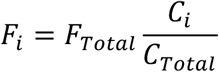

where *F_i_* is the research focus on resistance to drug class *i, F_Total_* is the total research activity (e.g. grants, publications, etc.) across all drug classes, *C_i_* is the consumption level for drug class *i*, and *C_Total_* is the total consumption across all drug classes. We again log_10_ transformed all data prior to analysis. We used linear models to test for deviations from expected research focus based on proportionality to drug usage (i.e. expectations based on proportion of total research attention equalling proportion of total antibiotic consumption), and found in all instances that this relationship significantly deviated from proportionality (Supplementary Fig. 7, Supplementary Table 1). In addition, as more newly discovered classes of antimicrobials could conceivably receive more research attention owing to less being known about their resistance mechanisms we also ran linear models assessing the relationship between log_10_ transformed measures of research focus and the number of years since discovery of an antimicrobial class. In all cases this relationship was non-significant (Supplementary Fig. 8, Supplementary Table 2).

### Data availability

The data used to analyse AMR research focus that support the findings of this study are provided with the paper as supplementary information (AMR research focus source data). The ECDC European antimicrobial resistance data are publicly available from the European Antimicrobial Resistance Surveillance Network Atlas (http://atlas.ecdc.europa.eu).

## References

1. Liu, Y.-Y. et al. Emergence of plasmid-mediated colistin resistance mechanism MCR-1 in animals and human beings in China: a microbiological and molecular biological study. Lancet Infect. Dis. 16, 161–168 (2016).

2. Blair, J. M. A., Webber, M. A., Baylay, A. J., Ogbolu, D. O. & Piddock, L. J. V. Molecular mechanisms of antibiotic resistance. Nat. Rev. Microbiol. 13, 42–51 (2015).

3. US Centers for Disease Control and Prevention. Antibiotic resistance threats in the United States, 2013. Centers for Disease Control and Prevention [online] (2013). Available at: http://www.cdc.gov/drugresistance/threat-report-2013/.

4. Surveillance Atlas of Infectious Diseases. Available at: http://atlas.ecdc.europa.eu/public/index.aspx. (Accessed: 27th November 2017)

5. Piddock, L. J. The crisis of no new antibiotics—what is the way forward? Lancet Infect. Dis. 12, 249–253 (2012).

6. Van Boeckel, T. P. et al. Global antibiotic consumption 2000 to 2010: an analysis of national pharmaceutical sales data. Lancet Infect. Dis. 14, 742–750 (2014).

7. Public Health Profiles. Available at: https://fingertips.phe.org.uk/profile/atlas-of-variation. (Accessed: 26th September 2017)

8. Antimicrobial consumption database (ESAC-Net). European Centre for Disease Prevention and Control Available at: http://ecdc.europa.eu/en/antimicrobial-consumption/surveillance-and-disease-data/database. (Accessed: 4th September 2017)

9. The Review on Antimicrobial Resistance. Tackling drug-resistant infections globally: final report and recommendations. (HM Government/Wellcome Trust, 2016) Available at: https://amr-review.org/. (Accessed: 4th September 2017)

10. Goff, D. A. et al. A global call from five countries to collaborate in antibiotic stewardship: united we succeed, divided we might fail. Lancet Infect. Dis. 17, e56–e63 (2017).

11. Köser, C. U., Ellington, M. J. & Peacock, S. J. Whole-genome sequencing to control antimicrobial resistance. Trends Genet. 30, 401–407 (2014).

12. Burnham, C.-A. D., Leeds, J., Nordmann, P., O’Grady, J. & Patel, J. Diagnosing antimicrobial resistance. Nat. Rev. Microbiol. 15, 697–703 (2017).

13. Toprak, E. et al. Evolutionary paths to antibiotic resistance under dynamically sustained drug selection. Nat. Genet. 44, 101–105 (2012).

14. Holmes, A. H. et al. Understanding the mechanisms and drivers of antimicrobial resistance. The Lancet 387, 176–187 (2016).

15. Vale, P. F. et al. Beyond killing: Can we find new ways to manage infection? Evol. Med. Public Health 2016, 148–157 (2016).

16. World Health Organisation. WHO publishes list of bacteria for which new antibiotics are urgently needed. World Health Organisation Available at: http://www.who.int/mediacentre/news/releases/2017/bacteria-antibiotics-needed/en/. (Accessed: 4th June 2017)

17. Targeting innovation in antibiotic drug discovery and development: the need for a One Health–One Europe–One World Framework (2016). (2017). Available at: http://www.euro.who.int/en/publications/abstracts/targeting-innovation-in-antibiotic-drug-discovery-and-development-the-need-for-a-one-healthone-europeone-world-framework-2016. (Accessed: 5th November 2018)

18. Czaplewski, L. et al. Alternatives to antibiotics—a pipeline portfolio review. Lancet Infect. Dis. 16, 239–251 (2016).

19. Tedijanto, C., Olesen, S., Grad, Y. & Lipsitch, M. Estimating the proportion of bystander selection for antibiotic resistance in the US. bioRxiv 288704 (2018). doi:10.1101/288704

20. Costelloe, C., Metcalfe, C., Lovering, A., Mant, D. & Hay, A. D. Effect of antibiotic prescribing in primary care on antimicrobial resistance in individual patients: systematic review and meta-analysis. BMJ 340, c2096 (2010).

21. Goossens, H., Ferech, M., Stichele, R. V. & Elseviers, M. Outpatient antibiotic use in Europe and association with resistance: a cross-national database study. The Lancet 365, 579–587 (2005).

22. Read, A. F. & Huijben, S. PERSPECTIVE: Evolutionary biology and the avoidance of antimicrobial resistance. Evol. Appl. 2, 40–51 (2009).

23. Abed, N. et al. An efficient system for intracellular delivery of beta-lactam antibiotics to overcome bacterial resistance. Sci. Rep. 5, srep13500 (2015).

24. Bikard, D. et al. Exploiting CRISPR-Cas nucleases to produce sequence-specific antimicrobials. Nat. Biotechnol. 32, 1146–1150 (2014).

25. Huang, J. M.-Y. et al. Rapid Electrochemical Detection of New Delhi Metallo-beta-lactamase Genes To Enable Point-of-Care Testing of Carbapenem-Resistant Enterobacteriaceae. Anal. Chem. 87, 7738–7745 (2015).

26. Raman, G., Avendano, E., Berger, S. & Menon, V. Appropriate initial antibiotic therapy in hospitalized patients with gram-negative infections: systematic review and meta-analysis. BMC Infect. Dis. 15, 395 (2015).

27. McAdams, D., Waldetoft, K. W., Tedijanto, C., Lipsitch, M. & Brown, S. P. Resistance diagnostics as a public health tool to combat antibiotic resistance: A model-based evaluation. bioRxiv 452656 (2018). doi:10.1101/452656

28. NIH Research Portfolio Online Reporting Tools (RePORT). Available at: https://report.nih.gov/index.aspx. (Accessed: 27th November 2017)

29. Altmetric. Altmetric Available at: https://www.altmetric.com/. (Accessed: 26th September 2017)

